# Insight towards the effect of the multibasic cleavage site of SARS-CoV-2 spike protein on cellular proteases

**DOI:** 10.1101/2020.04.25.061507

**Authors:** Kamal Shokeen, Shambhavi Pandey, Manisha Shah, Sachin Kumar

## Abstract

Severe respiratory syndrome coronavirus 2 (SARS-CoV-2) infection presents an immense global health problem. Spike (S) protein of coronavirus is the primary determinant of its entry into the host as it consists of both receptor binding and fusion domain. While tissue tropism, host range, and pathogenesis of coronavirus are primarily controlled by the interaction of S protein with the cell receptor, it is possible that proteolytic activation of S protein by host cell proteases also plays a decisive role. The host-cell proteases have shown to be involved in the proteolysis of S protein and cleaving it into two functional subunits, S1 and S2, during the maturation process. In the present study, the interaction of S protein of SARS-CoV-2 with different host proteases like furin, cathepsin B, and plasmin has been analyzed. Incorporation of the furin cleavage site (R-R-A-R) in the S protein in SARS-CoV-2 has been studied by mutating the individual amino acid. Our results suggest the polytropic nature of the S protein of SARS-CoV-2. Our analysis indicated that a single amino acid substitution in the polybasic cleavage site of S protein perturb the binding of cellular proteases. This mutation study might help to generate an attenuated SARS-CoV-2. Besides, targeting of host proteases by inhibitors may result in a practical approach to stop the cellular spread of SARS-CoV-2 and to develop its antiviral.

## Introduction

Coronaviruses (CoVs) possesses a single-stranded positive-sense RNA genome ranging from 26-32 kilobases in length (Weiss and Navas-Martin, 2005). The subfamily *Coronavirinae* contains a significant number of avian and mammalian pathogens. The subfamily *Coronavirinae* include Alphacoronavirus, Betacoronavirus, Gammacoronavirus, and Deltacoronavirus. To date, six different CoV strains are known to infect humans. The CoVs infecting humans belong to genera alphacoronavirus and betacoronavirus. The alphacoronaviruses infecting humans are HCoV-229E and HCoV-NL63, while betacoronaviruses infecting humans are HCoV-HKU1, HCoV-OC43, Middle East respiratory syndrome (MERS-CoV) and severe acute respiratory syndrome coronavirus (SARS-CoV).

The recent pandemic of severe respiratory syndrome coronavirus 2 (SARS-CoV-2) also belongs to betacoronavirus. All CoVs share correlation in their genome organization. It consists of 16 non-structural proteins (nsp1 to nsp16), encoded by open reading frame (ORF) 1a/b at the 5’ end, followed by the structural proteins spike (S), envelope (E), membrane (M), and nucleocapsid (N), encoded by other ORFs at the 3’ end. The entry of coronavirus into the host is a compounded process of receptor binding and proteolytic cleavage of S protein into functional subunits to promote virus-host cell membrane fusion (Heald-Sargent and Gallagher, 2012). The entry can be facilitated by employing fusion either directly at the cell surface receptor or through the endosomal compartment (Millet and Whittaker, 2015). The protruding S protein is responsible for the attachment of the virion to the cell surface receptor and its fusion, enabling the release of the viral genome into the cytoplasm (Millet and Whittaker, 2015). The ectodomain of the S protein consists of two functional subunits, S1 and S2. The S1 subunit is functionally responsible for receptor binding, while the S2 subunit maintains the fusion machinery process (Millet and Whittaker, 2015). The S protein is a class I viral fusion protein (Bosch et al., 2003), and is activated by its proteolytic cleavage by host-cell proteases (White et al., 2008). A variety of host-cell proteases like cathepsin B, trypsin, plasmin, elastase, and cell surface transmembrane protease/serine (TMPRSS) have been shown to cleave the S protein of SARS-CoVs to facilitate the viral attachment (Belouzard et al., 2012). It has been shown that the processing of surface glycoproteins is pre-requisite to produce comparatively more pathogenic and virulent viral particles and inhibition of these proteases likely to inhibit the viral infection as well (Zhou et al., 2015). Proteolytic activation unlocks the fusogenic potential of viral envelope glycoproteins and it is a crucial step in the entry of enveloped virus.

Recent studies have reported the incorporation of the unique furin cleavage site of Arg-Arg-Ala-Arg (R-R-A-R) between the boundary of S1 and S2 subunits in the S protein in SARS-CoV-2 (Coutard et al., 2020). Furin is the calcium-dependent protease that cleaves in an acidic environment after recognizing the specific sequence motif comprises of R-X-R/K-R, where X is any amino acid residue (Izaguirre, 2019). The sequence motifs generally present at the S1/S2 junction of S protein determines the cleavage by many host-cell proteases. Coronavirus has evolved for the proteolytic activation of S protein, and a large number of host proteases can have cognate recognition domain for its cleavage.

In the present study, we showed the binding of host-cell proteases (furin, cathepsin B, and plasmin) to the S protein of SARS-CoV-2. Furthermore, we analyzed the role of individual residues in the binding of S protein with host-cell proteases by incorporating the single point mutations in its cleavage site.

## Materials and methods

### Sequence analysis of S protein and its structure modeling

The SARS-CoV-2 sequences were retrieved from GenBank, accession number of each sequence used for the analysis is enlisted in (Supplementary Table S1). The sequences were aligned using the ClustalW algorithm implemented in the sequence alignment program package of MEGA-X software. Three-dimensional model of S protein was generated by using the SWISS-MODEL protein structure homology-modeling server (Waterhouse et al., 2018), wherein SARS-CoV-2 S glycoprotein (PDB id: 6VSB) was taken as a template. The energy of the modeled structure was minimized by using Yasara software, and finally, validated using PROCHECK (https://doi.org/10.1107/S0021889892009944). All the three-dimensional structures were visualized using PyMOL.

### Structure modeling of S protein mutants

Four different mutants of S protein were constructed with a single residue substitution in amino acid sequence. The mutations were incorporated with proline-to-alanine (P681A), arginine-to-alanine (R682A), arginine-to-alanine (R683A), and arginine-to-alanine (R685A). Further, the three-dimensional structures of the mutants were generated by SWISS-MODEL protein structure homology-modeling and further validated by PROCHECK-Ramachandran plot after its energy minimization using Yasara software.

### Molecular docking

The crystal structure for human plasminogen (PDB id 1DDJ), furin (PDB id 5MIM), and cathepsin (PDB id 1PBH) were retrieved from the protein data bank. The interaction between the proteases and SARS-CoV-2 S protein was performed by molecular docking. Docking between receptor (proteases) and ligand (S protein) was performed using Cluspro 2.0 protein-protein docking software (Kozakov et al., 2013, Kozakov et al., 2017, Vajda et al., 2017). For furin and cathepsin B, the C-chain of S protein, and for plasmin, the A-chain of S protein was used for the docking studies. The PDB sum generator server was used to find the interacting amino acid residues spanning the domain of proteases and S protein of SARS-CoV-2 (de Beer et al., 2014). Finally, the docked models were analyzed for Z-score using the DockScore tool (Malhotra et al., 2015).

## Results

The alignment of S protein amino acid sequences of SARS-CoV-2 with other CoVs provided critical insights into a unique mutation incorporated in its S1-S2 cleavage site (Figure 1). The unique incorporation of 4 amino acid residues, P681, R682, R683, and A684, were exclusive to the cleavage site of S protein of SARS-CoV-2.

**Figure 1.**
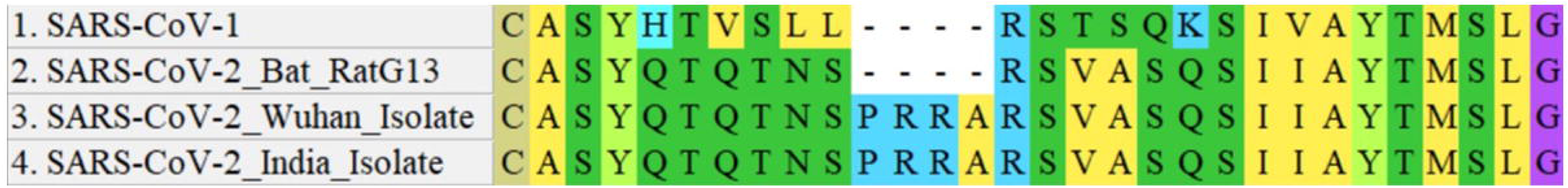
Sequence alignment of S protein of SARS-CoV isolates. Sequence alignment was analyzed using the S protein of SARS-CoV1, SARS-CoV-2, Bat RATG13 Isolate, SARS-CoV-2 India Isolate, and SARS-CoV-2 Wuhan Isolate.

The three-dimensional homo-trimer model of SARS-CoV-2 S protein showed 86.3% residues in most favoured regions, 12.6% in additional allowed regions, 0.6% in generously allowed regions and only 0.5% residues in disallowed regions (Figure 2). The final energy after the minimization of the model selected for further analysis was found to be -1715047.7 kJ/mol. The wild type S protein showed a significant interaction with furin protease (Figure 3A and 3F) with a total of 28 residues of S protein and 27 residues of furin involved in the binding. The interaction includes seven salt bridges, 13 hydrogen bonds, and 198 non-bonded contacts (Table 1). Furthermore, the interactive models of furin with each of the mutant S protein along with their interacting residues are shown in figure 3. The interaction studies of furin protease with different mutants of S protein revealed all the residues participated in hydrogen bond formation and salt bridges in the binding complex (Supplementary Table S2 and S3).

**Table 1.**
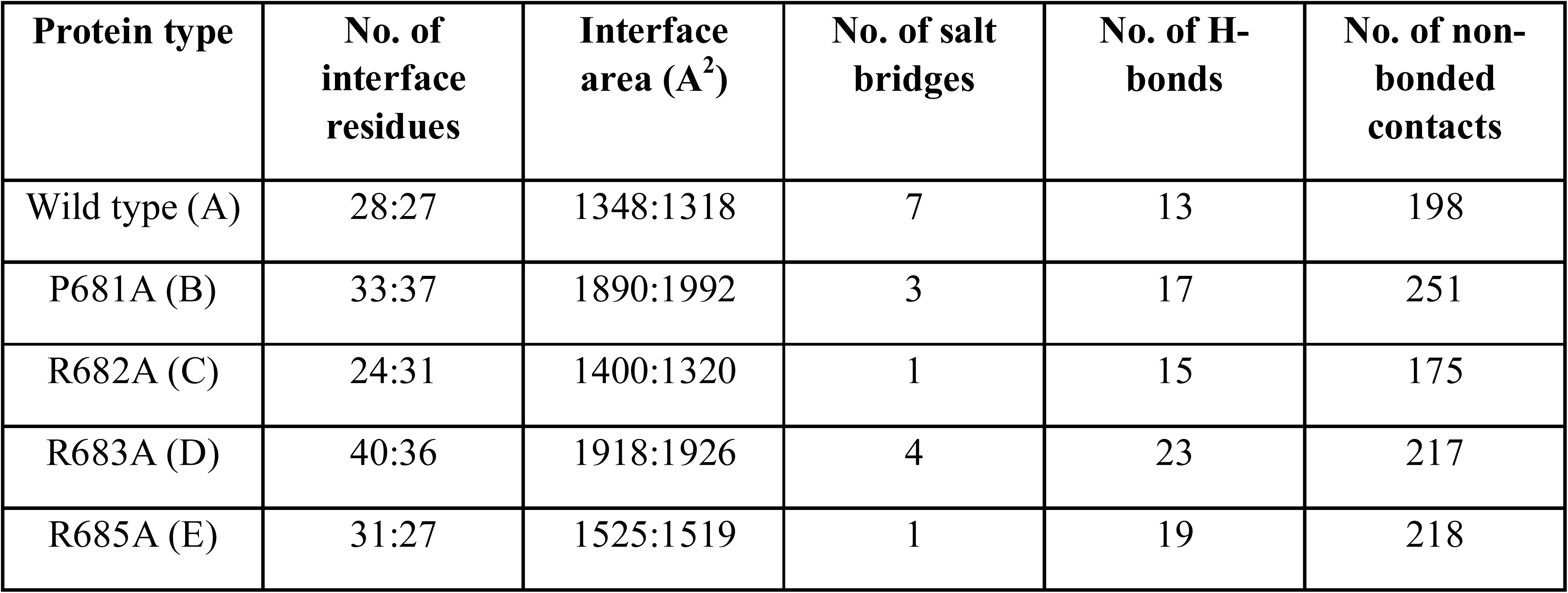
Representing the interface statistics of binding complex of furin enzyme with wild type S protein (A), P681A mutants (B), R682A mutants (C), R683A mutants (D), and R685A mutants (E).

**Figure 2.**
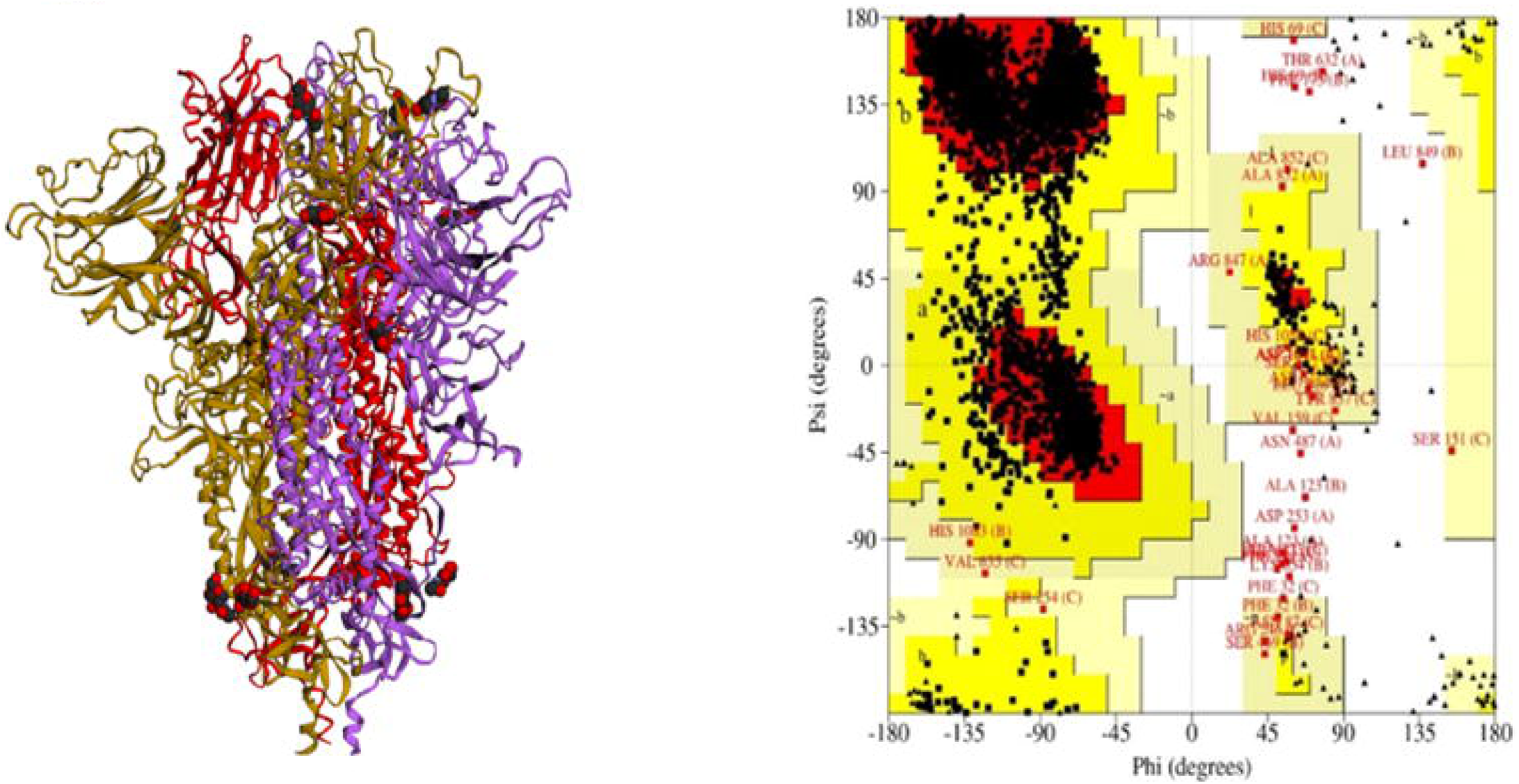
Structural details of S protein of SARS-CoV. The three-dimensional homo trimer model of SARS-CoV-2 S protein was analyzed using Ramachandran plot.

**Figure 3.**
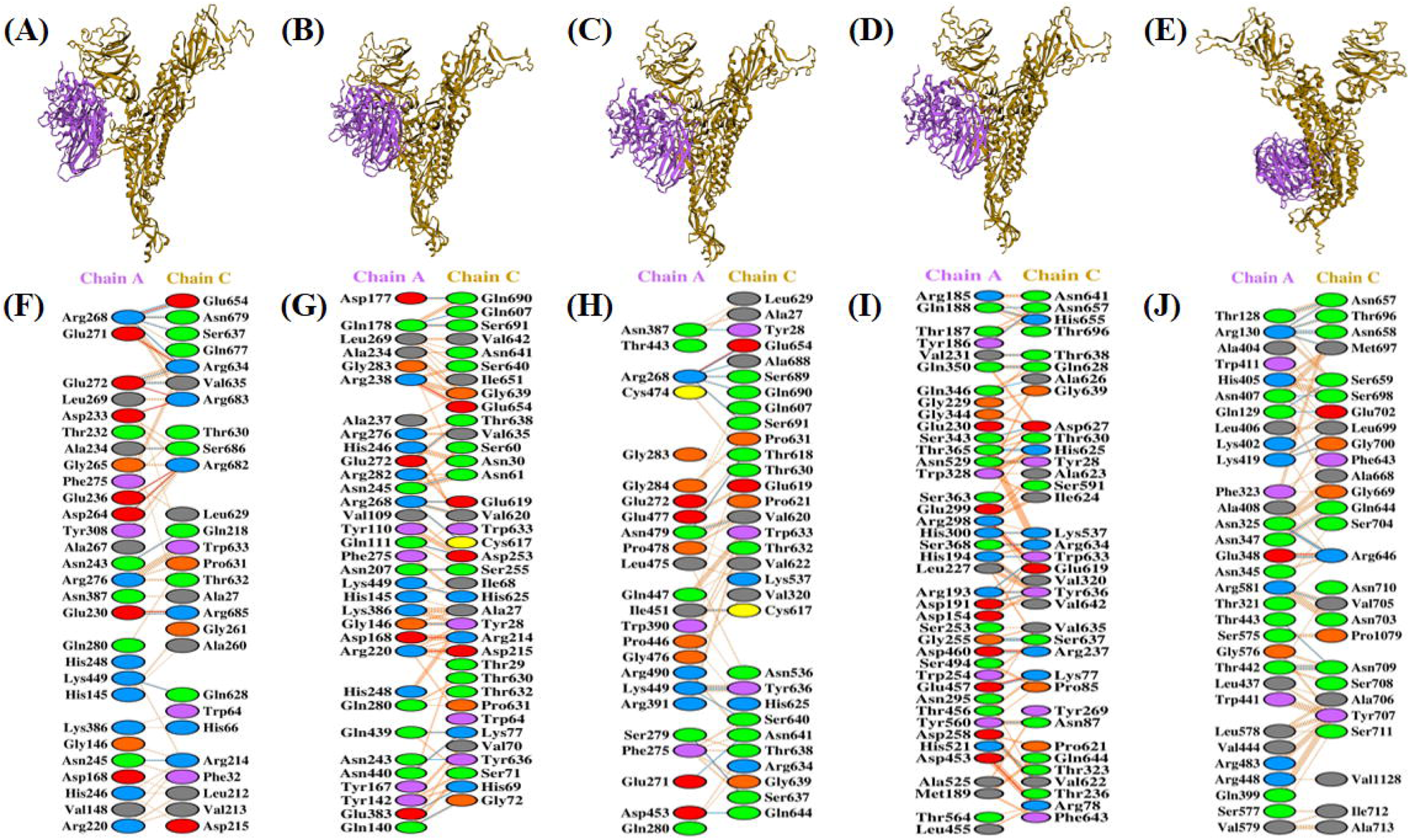
Visualization of docked complexes and interactive residues. The 3D-models of enzyme-protein binding complexes and their interacting amino acid residues. The 3D-binding complexes of A-chain of enzyme furin (violet) and C-chain of wild type S protein (golden) (A), P681A mutant (B), R682A mutant (C), R683A mutant (D), and R685A mutant (E). Interacting amino acid residues of A-chain of furin proteases and C-chain of wild type S protein (F), P681A mutant (G), R682A mutant (H), R683A mutant (I), and R685A mutant (J).

Similarly, the docking studies between the S protein and cathepsin B protease provided all the interaction details, including hydrogen bond and salt bridges forming residues (Figure 4). The final model evinced six salt bridges, 22 hydrogen bonds, and a total of 270 non-bonded contacts among wild type S protein and cathepsin B protease (Table 2). All the interacting residues for each model are enlisted in supplementary table S2 and S3, respectively. Docking studies between wild type and mutants of S protein with plasmin suggested interacting residues, including both hydrogen bond and salt bridges (Figure 5). The total of 31 residues of wild type S protein showed interaction with 38 residues of plasmin that includes 5 salt bridges, 18 hydrogen bonds and 275 non-bonded contacts (Table 3). Besides, the interaction of mutant S protein with plasmin showed interacting residues, including hydrogen bonds and salt bridges. Among the four mutants of S protein, the mutant with the least stability was selected based on the number of salt bridge present. The docking result of mutant with R685A substituted S protein with plasmin showed 1 salt bridges, 22 hydrogen bonds and 200 non-bonded contacts.

**Table 2.** Signifying the interface statistics of the binding complex of cathepsin B enzyme with wild type S protein (A), P681A mutants (B), R682A mutants (C), R683A mutants (D), and R685A mutants (E).

| **Protein type** | **No. of interface residues** | **Interface area (Å<sup>2</sup>)** | **No. of salt bridges** | **No. of H-bonds** | **No. of non-bonded contacts** |
| --- | --- | --- | --- | --- | --- |
| Wild type (A) | 37:29 | 1601:1661 | 6 | 22 | 270 |
| P681A (B) | 31:23 | 1333:1411 | 1 | 8 | 149 |
| R682A (C) | 43:31 | 1640:1664 | 1 | 12 | 224 |
| R683A (D) | 25:31 | 1519:1406 | 3 | 16 | 188 |
| R685A (E) | 36:33 | 1744:1759 | 2 | 15 | 182 |

**Table 3:** Table representing the interface statistics of the binding complex of plasmin enzyme with wild type S protein (A), P681A mutants (B), R682A mutants (C), R683A mutants (D), and R685A mutants (E).

| **Protein type** | **No. of interface residues** | **Interface area (Å<sup>2</sup>)** | **No. of salt bridges** | **No. of H-bonds** | **No. of non-bonded contacts** |
| --- | --- | --- | --- | --- | --- |
| Wild type (A) | 31:38 | 1857:1755 | 5 | 18 | 275 |
| P681A (B) | 42:46 | 1962:2026 | 5 | 22 | 329 |
| R682A (C) | 35:32 | 1599:1702 | 3 | 19 | 213 |
| R683A (D) | 33:30 | 1497:1479 | 4 | 17 | 179 |
| R685A (E) | 33:32 | 1620:1764 | 1 | 22 | 200 |

**Figure 4.**
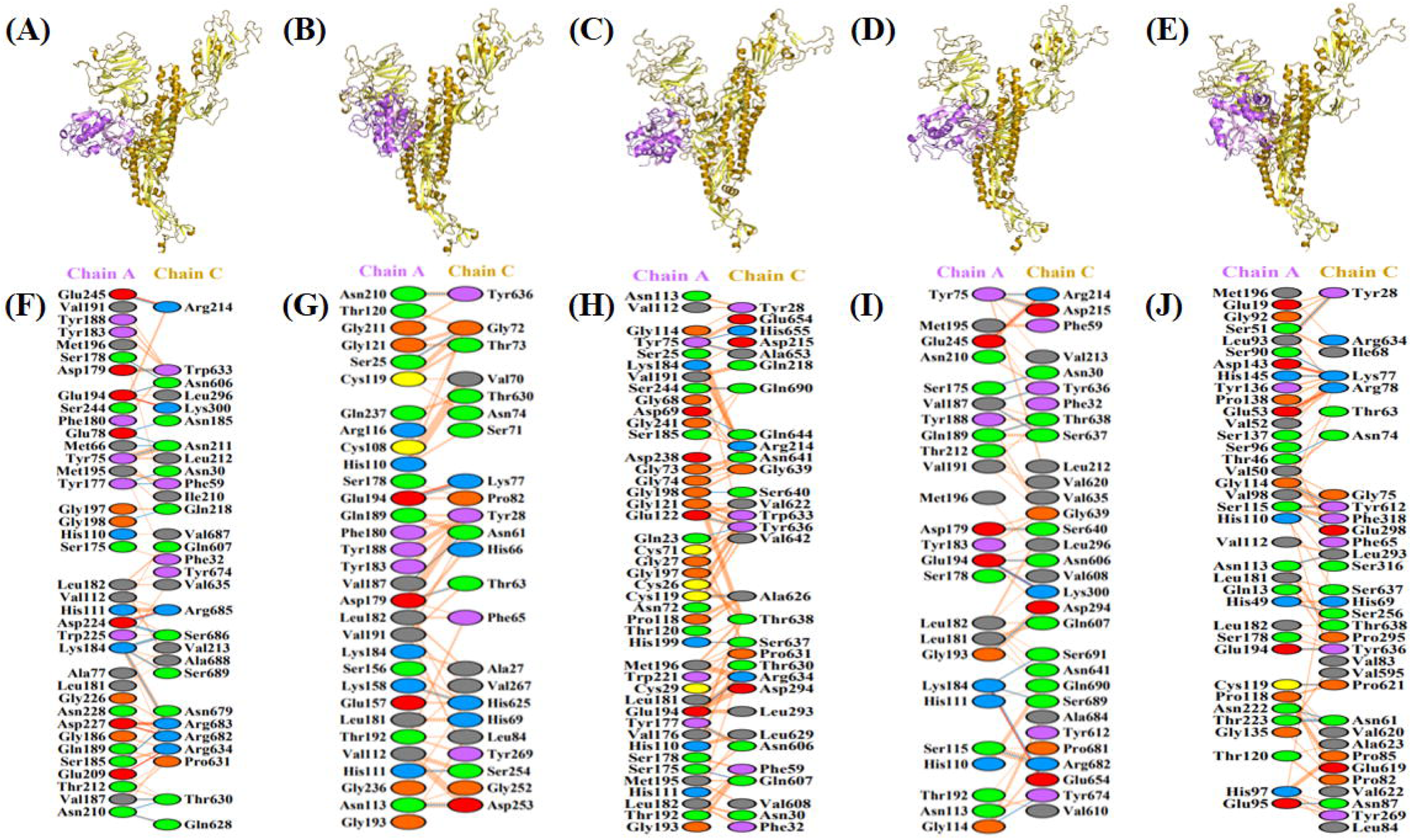
Visualization of docked complexes and interactive residues. The 3D-models of enzyme-protein binding complexes and their interacting amino acid residues. The 3D-binding complexes of A-chain of enzyme cathepsin B (violet) and C-chain of wild type S protein (golden) (A), P681A mutant (B), R682A mutant (C), R683A mutant (D), and R685A mutant (E). Interacting amino acid residues of A-chain of cathepsin B proteases and C-chain of wild type S protein (F), P681A mutant (G), R682A mutant (H), R683A mutant (I), and R685A mutant (J).

**Figure 5.**
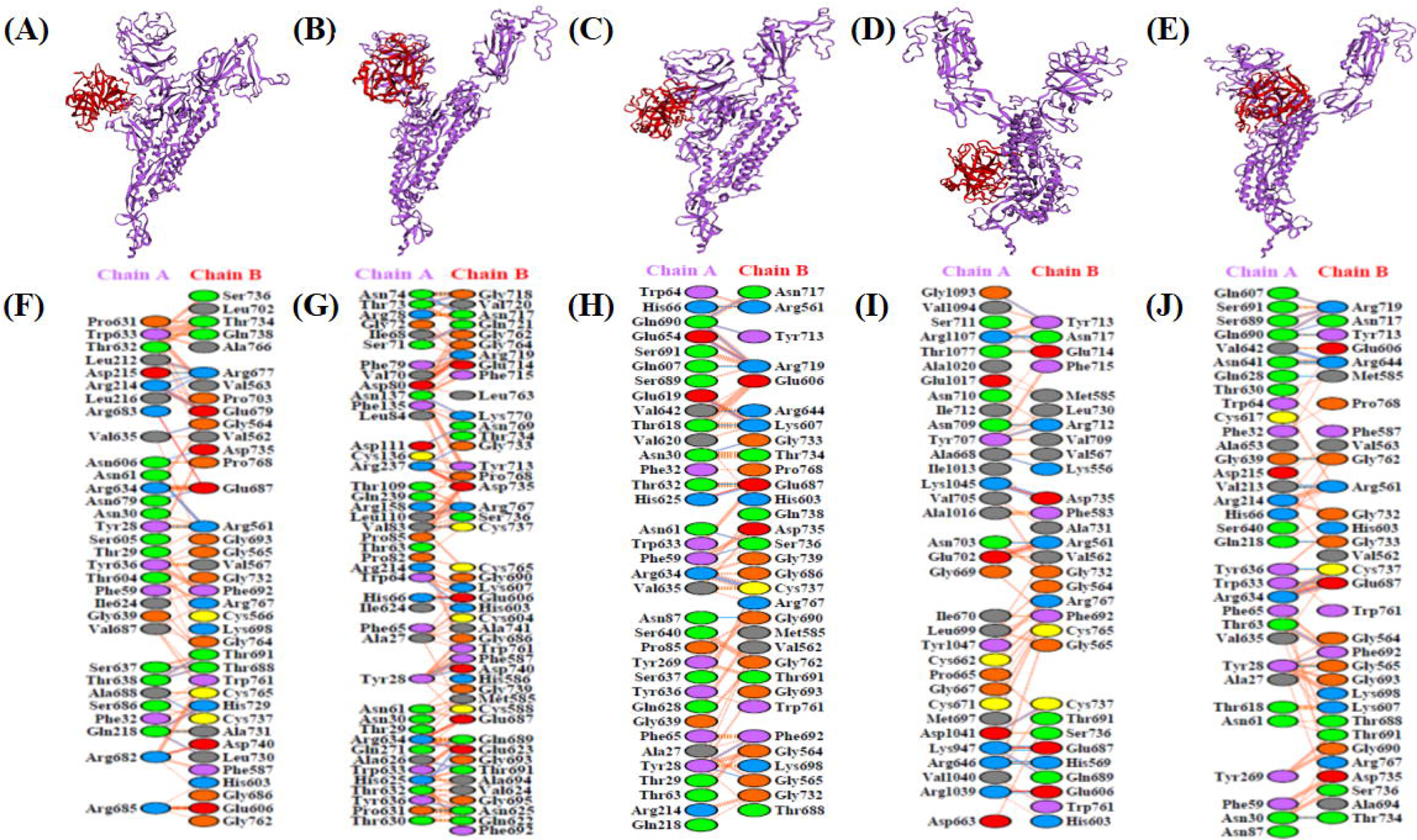
Visualization of docked complexes and interactive residues. The 3D-models of enzyme-protein binding complexes and their interacting amino acid residues. The 3D-binding complexes of B-chain of enzyme plasmin (Red) and A-chain of wild type S protein (violet) (A), P681A mutant (B), R682A mutant (C), R683A mutant (D), and R685A mutant (E). Interacting amino acid residues of B-chain of enzyme plasmin and A-chain of wild type S protein (F), P681A mutant (G), R682A mutant (H), R683A mutant (I), and R685A mutant (J).

## Discussion

It is known that the sequence motifs present between the boundary of S1 and S2 subunits of S protein in SARS-CoV determines the active binding and cleavage site for host-cell proteases (White et al., 2008). The results of the multiple sequence alignment suggested the incorporation of four additional polybasic amino acid residues at the S1/S2 site, as reported previously (Coutard et al., 2020). These amino acids signify the involvement of furin as host protease in the proteolytic processing of S protein. The presence of the polybasic cleavage sites for host-cell proteases is shown to directly impact the viral pathogenicity and host range (Nao et al., 2017). Several other viruses, including zika virus (Nambala and Su, 2018), dengue virus (Yu et al., 2008), avian influenza virus (Alexander and Brown, 2009), and Newcastle disease virus (Kumar and Kumar, 2014, Mohamed et al., 2011) have also been shown to modulate their pathogenicity by acquiring basic amino acids containing cleavage sites. The docking results of different host-cell proteases revealed their binding to the S protein through the formation of salt bridges and hydrogen bond linkages. The analysis of the residues involved in the interaction shows their presence in the binding complex. To emphasize the role of these additional amino acid residues in the binding of host-cell proteases to the S protein, we sequentially mutated the residues basic amino acid to alanine. Further, we analyzed the binding efficiency of each of the mutant with respective host protease by molecular docking. Firstly, we determined all the residues involved in the binding of host-cell proteases, including furin, cathepsin B, and plasmin with wild-type S protein. The proteomic identification of protease cleavage sites profiling of cathepsin B has revealed a strong cleavage site specificity for amino acid residue glycine and a partial preference for phenylalanine (Biniossek et al., 2011). The action of pH-independent cysteine protease cathepsin B is vital for the entry, hence the establishment of infection of SARS-CoV (Gierer et al., 2013). Our docking study denotes crucial details regarding the interaction and binding of cathepsin at the proteolytic cleavage site of SARS-CoV-2 S protein. The enzyme-protein interaction is sufficiently compromised when a single residue is mutated at the cleavage site of the protein. The dwindled and reduced number of salt bridge and hydrogen bond interactions in all mutant models from the wild type model signify a weaker enzyme-protein binding at the mutated cleavage site. Mutation of P682A in the wild type S protein results in the best mutant model that blocks the enzyme active site for cellular proteases, owing to a minimum number of salt bridge and hydrogen-bond formation.

The four newly incorporated residues in SARS-CoV-2 have been ascertained to have an implication on the pathogenicity of the virus. The docking studies convey the functional inference of these incorporated residues on the binding of proteases to the cleavage site of S protein. On adequately mutating one amino acid residue at the cleavage site of S protein, the binding of the protein is sufficiently hindered, and the residues of this site are not involved in the interaction with the proteases. The proteolytic cleavage, hence activation of S protein is amply controlled. Plasmin is a crucial enzyme in fibrinolysis and its natural substrate are fibrinogen and fibrin. The cleavage of influenza virus by plasmin is well characterized (Berri et al., 2013, Goto et al., 2001, LeBouder et al., 2010). Role of plasmin has been studied in A/WSN/1993 H1N1 influenza virus where the hemagglutinin (HA) cleavage site of the virus governs the spread of infection in plasmin dependent manner (Sun et al., 2010). The pivotal role of plasmin in pathogenicity of influenza virus was explained by the distribution of mini-plasmin and plasmin fragments in epithelial cells of bronchioles (Murakami et al., 2001). The role of plasmin has also been demonstrated in fusion of respiratory syncytial virus (Dubovi et al., 1983). The SARS-CoV-2 is characterized by hyperfibrinolysis, as evident by high levels of D-dimers, a breakdown product of fibrinolysis; however, it has been reported that plasmin can cleave S protein of SARS-CoV in vitro. (Chen et al., 2020, Guan et al., 2020, Huang et al., 2020, Wang et al., 2020). The cleavage of new furin site in the S protein of SARS-CoV-2 by host proteases like plasmin may enhance its infectivity and spread in respiratory cells. The administration of antiproteases to suppress the activity of proteases in the respiratory system may prevent or diminish the entry of virus in the respiratory system.

All viral fusion protein undergoes a structural transition and finally attain a compact low energy structure. These conformational changes brought viral and cellular host membrane in close proximity, which induces their fusion, followed by the formation of the pore that allows viral genetic material to enter the cell (White et al., 2008). This is exemplified by the HA protein of highly pathogenic avian influenza virus, where conversion of the monobasic site, cleaved by a trypsin-like protease to a polybasic site, allow cleavage by ubiquitously expressed furin-like proteases facilitate the spread of the virus and making it more virulent (Klenk and Garten, 1994, Lazarowitz and Choppin, 1975). Our results of substitution mutation R682A of S protein showed the least interaction with furin protease compared to its wild type homolog. Also, substitution mutation P681A of S exhibits the least interaction with cathepsin similarly the substitution mutation of R685A of S displays minimum interaction with plasmin protease. The finding suggested that PRRARS amino acid motif in the type S protein are responsible for its proper binding with furin, cathepsin and plasmin host proteases and mutation of these residues impair its interaction. Endoproteolytic cleavage, usually at arginine, is a common post-translational modification for activating several proteins such as peptide hormones and growth factor (Klenk and Garten, 1994). Docking studies revealed that modifications at these particular sites might weaken the interaction between the S protein and the host-cell proteases. Interestingly, the P681 did not directly involve in the binding of any protease. Still, on mutating the residue with alanine (P681A), the binding of all the proteases hindered to a greater extent. This might be due to the fact that proline provides stiffness to the molecular structure, which is required to maintain a proper conformation. Perhaps replacing proline with alanine might result in a local conformation change that further prevents the binding of host-cell proteases to their specific cleavage site.

These single amino acid substitutions in the cleavage site helped in understanding the role of individual residue in the binding complex. The participation of these crucial amino acids located at the boundary of S1 and S2 subunits in the proteolytic processing step will provide a unique opportunity to develop a lower pathogenic strain of SARS-CoV-2, which can further be used for vaccine development studies. Considering the fact proven by this in silico analysis that the single amino acid substitutions can help to make an attenuated form of virus, further in vitro work is needed to validate the fact.

## Acknowledgments

The virus research in our lab is currently supported by the Department of Biotechnology, Government of India (BT/562/NE/U-Excel/2016, and BT/PR24308/NER/95/644/2017).

## References

Alexander DJ, Brown IH. History of highly pathogenic avian influenza. Rev Sci Tech. 2009;28:19–38.

Belouzard S, Millet JK, Licitra BN, Whittaker GR. Mechanisms of coronavirus cell entry mediated by the viral spike protein. Viruses. 2012;4:1011–33.

Berri F, Rimmelzwaan GF, Hanss M, Albina E, Foucault-Grunenwald ML, Le VB, et al. Plasminogen controls inflammation and pathogenesis of influenza virus infections via fibrinolysis. PLoS Pathog. 2013;9:e1003229.

Biniossek ML, Nagler DK, Becker-Pauly C, Schilling O. Proteomic identification of protease cleavage sites characterizes prime and non-prime specificity of cysteine cathepsins B, L, and S. J Proteome Res. 2011;10:5363–73.

Bosch BJ, van der Zee R, de Haan CA, Rottier PJ. The coronavirus spike protein is a class I virus fusion protein: structural and functional characterization of the fusion core complex. J Virol. 2003;77:8801–11.

Chen N, Zhou M, Dong X, Qu J, Gong F, Han Y, et al. Epidemiological and clinical characteristics of 99 cases of 2019 novel coronavirus pneumonia in Wuhan, China: a descriptive study. Lancet. 2020;395:507–13.

Coutard B, Valle C, de Lamballerie X, Canard B, Seidah NG, Decroly E. The spike glycoprotein of the new coronavirus 2019-nCoV contains a furin-like cleavage site absent in CoV of the same clade. Antiviral Res. 2020;176:104742.

de Beer TA, Berka K, Thornton JM, Laskowski RA. PDBsum additions. Nucleic Acids Res. 2014;42:D292–6.

Dubovi EJ, Geratz JD, Tidwell RR. Enhancement of respiratory syncytial virus-induced cytopathology by trypsin, thrombin, and plasmin. Infect Immun. 1983;40:351–8.

Gierer S, Bertram S, Kaup F, Wrensch F, Heurich A, Kramer-Kuhl A, et al. The spike protein of the emerging betacoronavirus EMC uses a novel coronavirus receptor for entry, can be activated by TMPRSS2, and is targeted by neutralizing antibodies. J Virol. 2013;87:5502–11.

Goto H, Wells K, Takada A, Kawaoka Y. Plasminogen-binding activity of neuraminidase determines the pathogenicity of influenza A virus. J Virol. 2001;75:9297–301.

Guan WJ, Ni ZY, Hu Y, Liang WH, Ou CQ, He JX, et al. Clinical Characteristics of Coronavirus Disease 2019 in China. N Engl J Med. 2020.

Heald-Sargent T, Gallagher T. Ready, set, fuse! The coronavirus spike protein and acquisition of fusion competence. Viruses. 2012;4:557–80.

Huang C, Wang Y, Li X, Ren L, Zhao J, Hu Y, et al. Clinical features of patients infected with 2019 novel coronavirus in Wuhan, China. Lancet. 2020;395:497–506.

Izaguirre G. The Proteolytic Regulation of Virus Cell Entry by Furin and Other Proprotein Convertases. Viruses. 2019;11.

Klenk HD, Garten W. Host cell proteases controlling virus pathogenicity. Trends Microbiol. 1994;2:39–43.

Kozakov D, Beglov D, Bohnuud T, Mottarella SE, Xia B, Hall DR, et al. How good is automated protein docking? Proteins. 2013;81:2159–66.

Kozakov D, Hall DR, Xia B, Porter KA, Padhorny D, Yueh C, et al. The ClusPro web server for protein-protein docking. Nat Protoc. 2017;12:255–78.

Kumar CS, Kumar S. Species based synonymous codon usage in fusion protein gene of Newcastle disease virus. PLoS One. 2014;9:e114754.

Lazarowitz SG, Choppin PW. Enhancement of the infectivity of influenza A and B viruses by proteolytic cleavage of the hemagglutinin polypeptide. Virology. 1975;68:440–54.

LeBouder F, Lina B, Rimmelzwaan GF, Riteau B. Plasminogen promotes influenza A virus replication through an annexin 2-dependent pathway in the absence of neuraminidase. J Gen Virol. 2010;91:2753–61.

Malhotra S, Mathew OK, Sowdhamini R. DOCKSCORE: a webserver for ranking protein-protein docked poses. BMC Bioinformatics. 2015;16:127.

Millet JK, Whittaker GR. Host cell proteases: Critical determinants of coronavirus tropism and pathogenesis. Virus Res. 2015;202:120–34.

Mohamed MH, Kumar S, Paldurai A, Samal SK. Sequence analysis of fusion protein gene of Newcastle disease virus isolated from outbreaks in Egypt during 2006. Virol J. 2011;8:237.

Murakami M, Towatari T, Ohuchi M, Shiota M, Akao M, Okumura Y, et al. Mini-plasmin found in the epithelial cells of bronchioles triggers infection by broad-spectrum influenza A viruses and Sendai virus. Eur J Biochem. 2001;268:2847–55.

Nambala P, Su WC. Role of Zika Virus prM Protein in Viral Pathogenicity and Use in Vaccine Development. Front Microbiol. 2018;9:1797.

Nao N, Yamagishi J, Miyamoto H, Igarashi M, Manzoor R, Ohnuma A, et al. Genetic Predisposition To Acquire a Polybasic Cleavage Site for Highly Pathogenic Avian Influenza Virus Hemagglutinin. mBio. 2017;8.

Sun X, Tse LV, Ferguson AD, Whittaker GR. Modifications to the hemagglutinin cleavage site control the virulence of a neurotropic H1N1 influenza virus. J Virol. 2010;84:8683–90.

Vajda S, Yueh C, Beglov D, Bohnuud T, Mottarella SE, Xia B, et al. New additions to the ClusPro server motivated by CAPRI. Proteins. 2017;85:435–44.

Wang D, Hu B, Hu C, Zhu F, Liu X, Zhang J, et al. Clinical Characteristics of 138 Hospitalized Patients With 2019 Novel Coronavirus-Infected Pneumonia in Wuhan, China. Jama. 2020.

Waterhouse A, Bertoni M, Bienert S, Studer G, Tauriello G, Gumienny R, et al. SWISS-MODEL: homology modelling of protein structures and complexes. Nucleic Acids Res. 2018;46:W296–w303.

Weiss SR, Navas-Martin S. Coronavirus pathogenesis and the emerging pathogen severe acute respiratory syndrome coronavirus. Microbiol Mol Biol Rev. 2005;69:635–64.

White JM, Delos SE, Brecher M, Schornberg K. Structures and mechanisms of viral membrane fusion proteins: multiple variations on a common theme. Crit Rev Biochem Mol Biol. 2008;43:189–219.

Yu IM, Zhang W, Holdaway HA, Li L, Kostyuchenko VA, Chipman PR, et al. Structure of the immature dengue virus at low pH primes proteolytic maturation. Science. 2008;319:1834–7.

Zhou Y, Vedantham P, Lu K, Agudelo J, Carrion R, Jr., Nunneley JW, et al. Protease inhibitors targeting coronavirus and filovirus entry. Antiviral Res. 2015;116:76–84.

