## Supplementary table 1, 2, 3 for "Insight towards the effect of the multibasic cleavage site of SARS-CoV-2 spike protein on cellular proteases"

| **Spike protein sequences of SARS-CoV-2 Isolates** | **Accession Number** |
| --- | --- |
| SARS-CoV-1 | ABD72968.1 |
| SARS-CoV-2 Bat RATG13 Isolate | QHR63300.2 |
| SARS-CoV-2 Wuhan Isolate | QHU36864.1 |
| SARS-CoV-2 India Isolate | QIA98583.1 |

Table S1: Denoting the spike protein sequences of different isolates of SARS-CoV-2 with their accession number used for the study and analysis.

| **Protein type** | **Host-cell Protease** | **Amino Acid Residues of SARS-CoV-2 S protein** | **Amino Acid Residues of protease** |
| --- | --- | --- | --- |
| Wild type (A) | Furin | R214, Q218, Q628, R634, V635, S637, E654, N679, R685, S686 | E230, A234, N243, N245, R268, E272, L449 |
|  | Cathepsin B | N30, F59, N185, N211, R214, Q218, N606, Q628, T630, W633, R683, R685, S686, S689 | M66, Y75, E78, Y177, S178, D179, K184, S185, V187, E194, G198, N210, D224, E225 |
|  | Plasmin | Y28, N61, L212, R214, D215, Q218, N606, V635, S637, T638, N606, N679, A688 | R561, G564, R677, E679, E687, T688, T691, H729, L730, A731, D735, C737 |
| P681A (B) | Furin | Y28, N61, H69, G72, L77, D253, S255, Q607, C617, E619, V620, H625, Y636, T638, Q690, S691 | Q111, Q140, G146, D177, Q178, N207, N243, N245, R268, E272, E383, N439, N440, L449 |
|  | Cathepsin B | T63, N61, G72, K77, D253, S254, H625, Y636 | S25, H111, N113, K158, D179, V187, E194, N210 |
|  | Plasmin | Y28, W64, H66, S71, T73, N74, F79, V83, L110, D111, N137, R237, Q271, A626, H625, P631, T632, Y636 | F587, H603, E606, N625, Q689, T691, G693, N717, G718, R719, V720, Q721, G733, C737, D740, R767, N769, K770 |
| R682A (C) | Furin | Y28, C617, T618, V620, Y636, S637, S640, N641, Q644, E654, A688, S689, Q690, S691 | R268, E271, S279, N387, K449, I451, D453, N479 |
|  | Cathepsin B | N606, Q607, A626, R634, Y636, S637, S640, Q644, A653, H655, Q690 | Q23, D69, Y75, C119, E122, S175, Y177, E194, G198, H199, S244 |
|  | Plasmin | Y28, N30, F59, N61, H66, N87, T618, E619, H625, T632, R634, V642, E654, Q607, Q690 | R561, G565, K607, R644, E687, G690, F692, Y713, N717, R719, G733, S736, C737 |
| R683A (D) | Furin | Y28, K77, N87, K537, V622, H625, A626, D627, Q628, W633, R634, Y636, S637, T638, H655, N657 | Y186, Q188, D191, R193, H194, V231, G255, D258, H300, S343, Q346, Q350, T365, S368, E457, D460, N529, Y560 |
|  | Cathepsin B | N30, R214, D215, D294, K300, N606, Y636, S637, T638, S640, E654, Y674, R682, S689, S691 | Y75, G114, S115, S175, D179, K184, V187, Q189, G193, E194 |
|  | Plasmin | R646, A668, N703, Y707, N709, K947, R1039, D1041, K1045, T1077, G1093, R1107 | K556, R561, H569, E606, E687, Q689, R712, Y713, E714, S717, D735, C737 |
| R685A (E) | Furin | Q644, R646, N657, N658, S659, T696, M697, S698, L699, G700, E702, V705, N709 | T128, Q129, R130, N347, E348, N407, K419, T442, T443, R581 |
|  | Cathepsin B | Y28, N61, N74, K77, N87, S256, L293, E298, Y612, P621, Y636 | T46, H49, S90, L93, E95, N113, G114, S115, C119, H145, E194, N222, T223 |
|  | Plasmin | Y28, N30, T63, V213, Q218, Q607, Q628, W633, R634, Y636, G639, N641, V642, S689, Q690 | R561, G565, M 585, R644, E687, F692, Y713, N717, R719, G733, T734, C737, G762 |

Table S2. Representing amino acid residues involved in forming hydrogen bond contacts in the binding complexes of the enzyme (furin, cathepsin B and plasmin) with wild type S protein (A), P681A mutants (B), R682A mutants (C), R683A mutants (D), and R685A mutants (E).

| **Protein type** | **Host-cell Protease** | **Amino Acid Residues of SARS-CoV-2 S protein** | **Amino Acid Residues of protease** |
| --- | --- | --- | --- |
| Wild type (A) | Furin | R634, E654, R682, R683, R685 | E230, D233, E236, D264, R268, E271, E272 |
|  | Cathepsin B | R214, K300, R634, R682, R683, R685 | E194, E209, D224, D227, E245 |
|  | Plasmin | R214, R634, R682, R683, R685 | E606, E679, E687, D740 |
| P681A (B) | Furin | R214, D215, E654 | D168, R220, R238 |
|  | Cathepsin B | K77 | E194 |
|  | Plasmin | H66, D80, R237, R634 | E606, E623, E687, R719, D735 |
| R682A (C) | Furin | E654 | R268 |
|  | Cathepsin B | R634 | E194 |
|  | Plasmin | E619, H625, E654 | K607, E687, R719 |
| R683A (D) | Furin | K77, R78, R237, E619 | R298, D453, E457, D460 |
|  | Cathepsin B | R214, K300, E654 | K184, E194, E245 |
|  | Plasmin | E702, K947, R1039, K1045 | R561, E606, E687, D735 |
| R685A (E) | Furin | R646 | E348 |
|  | Cathepsin B | K77, R78 | E53, D143 |
|  | Plasmin | R634 | E687 |

Table S3: Representing amino acid residues involved in forming salt bridge interactions in the binding complexes of the enzyme (furin, cathepsin B and plasmin) with wild type S protein (A), P681A mutants (B), R682A mutants (C), R683A mutants (D), and R685A mutants (E).
